# Phosphorylation of the rod-tail hinge region of cingulin regulates its interaction with nonmuscle myosin-2B

**DOI:** 10.64898/2026.04.02.716052

**Authors:** Florian Rouaud, Annick Mutero-Maeda, Christian Borgo, Maria Ruzzene, Sandra Citi

**Author notes:** Corresponding address: Prof. Sandra Citi.

## Abstract

The tight junction (TJ) protein cingulin binds directly to nonmuscle myosin 2B (NM2B) through sequences in its C-terminal rod-tail region and recruits it to tight junctions (TJ) to control membrane cortex mechanics, epithelial morphogenesis and cingulin conformation. However, the minimal sequence required for cingulin-NM2B interaction and how this interaction is regulated is not known. Here we identify a 19-aminoacid sequence at the hinge between the cingulin rod and tail that is required for cingulin-NM2B interaction, and we investigate the role of phosphorylation of Ser residues within this region in regulating this interaction. Immunofluorescence microscopy localization of NM2B in cingulin-KO cells rescued with mutant cingulin constructs shows that phospho-mimetic but not dephospho-mimetic cingulin mutants inhibit NM2B recruitment to junctions and downstream regulation of cingulin conformation and TJ tortuosity, correlating with cingulin-NM2B interaction, as determined by GST pulldown analysis. In contrast, either phospo-or dephospho-mimetic mutants of Ser residues within the cingulin head domain do not affect either NM2B recruitment to TJ, or cingulin conformation and localization in cells, or TJ membrane tortuosity. Finally, Ser residues within the hinge display the consensus sequence for protein kinases CK1 and CK2, and, through in vitro phosphorylation, site mutation analysis and use of inhibitors, we identify a complex interplay between CGN phospho-sites, with a prominent negative role of Ser1162 phosphorylation in the regulation of cingulin-NM2B interaction. In summary, we show that cingulin-NM2B interaction is regulated by cingulin phosphorylation within the hinge and identify a potential role for CK1 and CK2 kinases in cingulin phosphorylation.

## INTRODUCTION

Tight junctions (TJs) are essential for the barrier function of epithelial and endothelial tissues. They consist of transmembrane proteins, such as claudins, occludins and JAMs, and a cytoplasmic plaque comprising scaffolding proteins, such as ZO proteins, and adaptor proteins such as cingulin family proteins, which are structurally and functionally connected to the actomyosin and microtubule cytoskeletons (reviewed in (Citi et al., 2024; Otani and Furuse, 2020; Van Itallie and Anderson, 2014)). The dynamic remodeling of the actomyosin cytoskeleton by Rho GTPases and their downstream effectors is essential for the regulation of TJ assembly, disassembly and function during development, health and disease (reviewed in (Buckley and Turner, 2018; Quiros and Nusrat, 2014; Varadarajan et al., 2019)). The microtubule cytoskeleton is important for the establishment and maintenance of epithelial apicobasal polarity and is also connected to the epithelial junctional complex and to TJs (reviewed in (Akhmanova and Kapitein, 2022; Musch, 2004; Yano et al., 2017)}.

To understand the regulation of TJs by the cytoskeleton, it is important to determine which TJ proteins bind to cytoskeletal proteins, and how their interaction is regulated. ZO-1 (Stevenson et al., 1986) and cingulin family proteins (Citi et al., 1988; Guillemot and Citi, 2006; Ohnishi et al., 2004) play an important role in connecting TJs to the cytoskeleton. ZO-1 functions as a scaffold for TJ membrane proteins through its N-terminal regions, whereas its C-terminus binds to cingulin (D’Atri et al., 2002; Umeda et al., 2004; Vasileva et al., 2022). Both ZO-1 and cingulin bind to actin filaments (Belardi et al., 2020; D’Atri and Citi, 2001; Fanning et al., 1998; Fanning et al., 2002; Rouaud et al., 2023; Van Itallie et al., 2009; Wittchen et al., 1999). Cingulin and paracingulin bind to nonmuscle myosins (NM2s) to recruit them to epithelial and endothelial junctions in vitro and in vivo (Citi et al., 1988; Cordenonsi et al., 1999a; Rouaud et al., 2023; Rouaud et al., 2025a). Thus, the connection of TJs to the actomyosin cytoskeleton consists of a module comprising ZO-1, cingulin, F-actin and NM2s. Recruitment of NM2B to TJs by cingulin promotes TJ assembly of ZO-1, TJ membrane tortuosity, and epithelial morphogenesis (Rouaud et al., 2023; Rouaud et al., 2024; Vasileva et al., 2022). Moreover, both cingulin and paracingulin bind to microtubules and associated proteins, to regulate microtubule organization and epithelial polarity (Flinois et al., 2024a; Flinois et al., 2024b; Mangan et al., 2016; Yano et al., 2013). However, little is known about the mechanisms regulating the interaction of ZO-1 and cingulin family proteins with the cytoskeleton. For example, despite the fact that several TJ proteins, including ZO-1, cingulin, claudins and occludin are phosphorylated (Citi and Denisenko, 1995; Cordenonsi et al., 1997; Dorfel et al., 2013; Sakakibara et al., 1996; Stevenson et al., 1989; Van Itallie et al., 2012; Yamamoto et al., 2008), it is not clear whether phosphorylation regulates the interaction of these proteins with the cytoskeleton (reviewed in (Reiche and Huber, 2020; Shigetomi and Ikenouchi, 2018; Van Itallie and Anderson, 2018)). In the case of cingulin, it was reported that phosphorylation in the head region regulates cingulin conformation and interaction with microtubules in vitro (Yano et al., 2018). Regarding the potential involvement of specific protein kinases in the phosphorylation of TJ proteins, protein kinases CK1 and CK2 were shown to phosphorylate occludin and tricellulin (Cordenonsi et al., 1999b; Dorfel et al., 2013; McKenzie et al., 2006; Raleigh et al., 2011), whereas purified cingulin was shown to be phosphorylated by adenosine monophosphate-activated protein kinase (AMPK) to promote an open conformation in vitro (Yano et al., 2013; Yano et al., 2018). On the other hand, we recently showed by immunofluorescence microscopy analysis of the distance between cingulin N-terminal and C-terminal ends that at junctions cingulin can be present either in an open (extended) or closed (folded) conformation, correlating with its ability to bind NM2B (Rouaud et al., 2024). Thus, one open question is whether phosphorylation in the head region of cingulin affects cingulin conformation in cells.

Another potential region of cingulin phosphorylation is the rod-tail region. The coiled-coil rod of cingulin shows the highest sequence homology to the rod of NM2 heavy chains, and, similarly to NM2s, is followed by a small globular tail at its C-terminus (Citi et al., 2000; Cordenonsi et al., 1999a). Since phosphorylation of the myosin heavy chains at the C-terminal end of the rod and in the non-helical tail controls many aspects of NM2 function, including conformation, self-interaction and filament assembly (Babkoff et al., 2021; Cote et al., 1981; Dulyaninova and Bresnick, 2013; Dulyaninova et al., 2007; Egelhoff et al., 1993; Kawamoto and Adelstein, 1988; Rosenberg and Ravid, 2006; Rosenberg et al., 2008; Straussman et al., 2007) we hypothesized that cingulin phosphorylation in the rod-tail hinge region may regulate its interaction with NM2, which occurs through the C-terminal regions of both cingulin and NM2 (Rouaud et al., 2023). To address the role of rod-tail versus head phosphorylation, here we use immunofluorescence microscopy and GST pulldown assays to investigate cingulin-NM2B interaction, cingulin conformation and TJ membrane tortuosity. Through the analysis of cingulin mutants we identify a 19-residue sequence in the rod-tail hinge region, that is required for cingulin interaction with NM2B, and we show that phosphorylation of Ser residues within this region negatively regulates interaction with NM2B and downstream cingulin conformation and TJ membrane tortuosity. In contrast, mutations of Ser residues within the head have no effect on junctional recruitment of NM2B, cingulin conformation in cells, cingulin-NM2B interaction and TJ membrane tortuosity. Furthermore, we provide evidence for a potential role of CK1 and CK2 in the phosphorylation-dependent interaction between cingulin and NM2B.

## RESULTS

### Phosphorylation of Ser residues within a 19-residue sequence in the rod-tail hinge region controls cingulin-dependent NM2B recruitment to junctions

MDCK cells KO for cingulin show loss of junctional NM2B labeling, and expression of full-length cingulin but not of cingulin with a truncation of its C-terminal 187 residues restores junctional NM2B, indicating that the C-terminal part of the rod and the tail are critical regions for cingulin-NM2B interaction (Fig. 1A-C) (Rouaud et al., 2023). To narrow down the sequence responsible for cingulin-NM2B interaction and investigate the role of phosphorylation, we used as an assay the junctional localization of NM2B in the background of CGN-KO cells rescued with WT and mutant constructs. Analysis of phospho-proteomic databases indicated that cingulin phosphorylation occurs at evolutionarily conserved residues within a short sequence within the rod-tail hinge, specifically at Ser 1144, 1149, 1151, 1155, 1162, 1163, 1169, 1174 and 1178, of the canis sequence (Fig. 1D, top), corresponding to Ser residues at positions 1156, 1161, 1163, 1167, 1175, 1176, 1182, 1187 and 1191 in the human sequence (Fig. 1D, bottom). All these Ser residues are in the globular tail region of cingulin (aminoacids 1149-1190 in the canis sequence), except for the first Ser (1144 in canis), which is in the rod.

**Figure 1.**
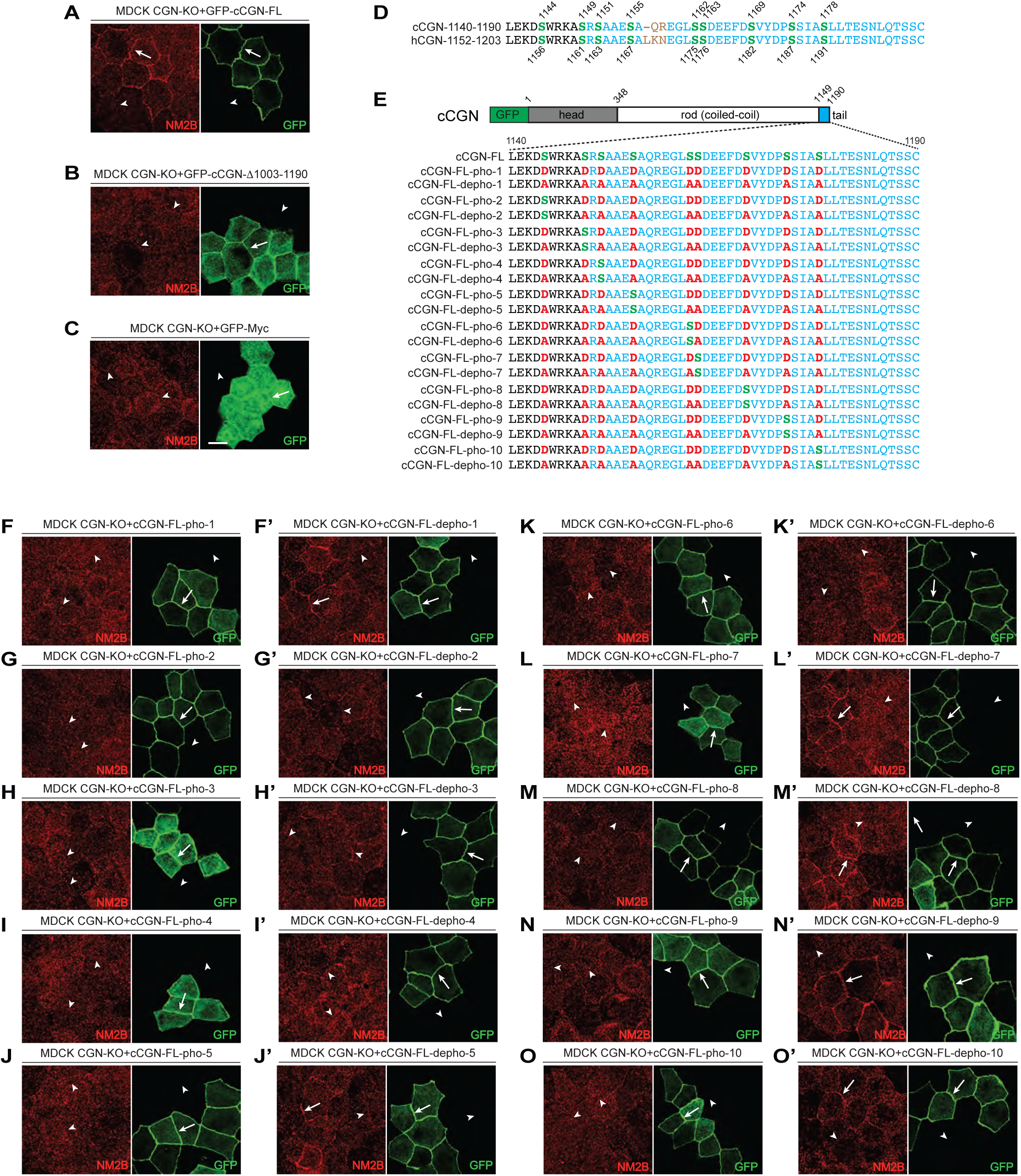
Phosphorylation of Ser residues in the rod-tail regions of cingulin inhibits the junctional accumulation of NM2B. (A-C) IF microscopy analysis and localization of junctional labeling of NM2B in CGN-KO MDCK cells rescued with GFP-cCGN-FL (A, positive control), or with GFP-cCGN-Δ1003-1190 (B), or with GFP–Myc alone (negative control, panel C). (D) Sequence alignment between the rod-tail hinge of the canis (top) and human (bottom) cingulin sequences, to highlight the conserved residues, specifically the Serines (in green). Residues in blue color belong to the tail region. (E) Top: scheme of GFP-tagged canine CGN (cCGN-FL), with the GFP tag (green), globular head (gray), coiled-coil rod (white) and globular tail (blue) domains (amino-acid residue boundaries are indicated above the scheme). Bottom: C-terminal sequences of WT canine CGN (cCGN-FL, region 1140–1190 with specific residues indicated in the region) and corresponding sequences of the cCGN-phospho-mimetic or dephospho-mimetic mutants. WT Ser residues are shown in green, and either phospho-(Asp-D) or dephospho (Ala-A) mutations are shown in red. Mutants pho-2/depho-2 to pho-10/depho-10 are based on pho-1 and depho-1, and contain only one mutated residue that is reverted from either D or A to the WT Ser. (F-O) IF microscopy analysis and localization of junctional labeling of NM2B in CGN-KO MDCK cells rescued with GFP-cCGN-pho-1 to pho-10 (F-O), or with GFP-cCGN-depho-1 to depho-10 (F’-O’). Arrows and arrowheads show normal and decreased/undetected junctional labeling, respectively. Scale bar: 10 μm.

Next, we mutated all the Ser residues in this region either to Asp (“cCGN-FL-pho-1” phospho-mimetic mutant, Fig. 1E) or to Ala (“cCGN-FL-depho-1” dephospho-mimetic mutant, Fig. 1E), and we analyzed their ability to rescue NM2B accumulation at TJs in the background of CGN-KO MDCK cells. Strikingly, the pho-1 mutant failed to rescue NM2B to junctions (arrowheads in Fig. 1F, arrows showing exogenous CGN pho-1 mutant in green), whereas the depho-1 mutant rescued NM2B, similar to WT cingulin (arrows in Fig. 1F’). This indicated that cingulin phosphorylation negatively regulates CGN-NM2B interaction at TJs. To identify more precisely the relevance of specific Ser residues, we generated 9 additional mutants, where starting from either pho-1 or depho-1, only one of each of the 9 mutated residues was reverted to Ser from either Asp or Ala, making it available, potentially, for phosphorylation (Fig. 1E, pho-2 to pho-10 and depho-2 to depho-10 mutants). Immunofluorescence microscopy analysis showed that none of the pho-mutants (pho-2 to pho-10) could rescue the junctional accumulation of NM2B (NM2B in red, arrowheads in Fig. 1G-O), suggesting that phosphorylation at redundant sites negatively regulates cingulin-NM2B interaction in cells, and no phosphorylation at any specific site overrides the negative effect of phosphorylation at other sites. Among the dephosphorylated mutants where only one Ser was changed back from Ala to Ser, depho-5, depho-7, depho-8, depho-9 and depho-10 rescued NM2B similarly to depho-1 (arrows in Fig. 1J’, L’, M’, N’ and O’, respectively), indicating that allowing phosphorylation at Ser 1167, 1176 1182, and 1191 does not negatively impact cingulin-NM2B interaction. In contrast, the depho-2, depho-3, depho-4 and depho-6 mutants failed to rescue junctional NM2B (arrowheads in Fig. 1G’, H’, I’ and K’, respectively), suggesting that allowing phosphorylation at either Ser1144, 1149, 1151 or 1162 has a negative impact on cingulin-NM2B interaction, even when all other residues cannot be phosphorylated. All the cingulin mutants were correctly recruited to junctions (Fig. 1F-O, arrows in green channel), consistent with the notion that binding to ZO-1 through the ZIM region in the cingulin head is the most critical factor controlling cingulin junctional recruitment (D’Atri et al., 2002) and indicating that the phosphorylation of the rod-tail does not affect cingulin-ZO-1 interaction.

Based on the above mutagenesis analysis, we focused our attention on the sequence 1144-1162, that comprises the residues whose phosphorylation is critical to control cingulin-NM2B interaction. This region comprises 5 residues of the coiled-coil rod (1144-1148 in the canis sequence) and 14 residues of the non-helical tail (1149-1162 in the canis sequence), thus constituting the 19-residue rod-tail “hinge” region (Fig. 2A, scheme of cingulin in the middle, alignment of sequences of WT and mutants top and bottom, with hinge region in brown). To analyze the role of the hinge region and phosphorylation of Ser residues within it, we compared the behavior of a C-terminal truncation of the C-terminus cingulin (Δ1003-1190, Fig. 2A, 2D, Fig. 1B) with that of either a deletion of the hinge (Δ1144-1162, Fig. 2A, 2E), or of mutants of the hinge, where the 4 Ser residues were mutated either to Asp (phospho, Fig. 2A, 2F) or to Ala (dephospho, Fig. 2A, 2G). In addition, since phosphorylation of three residues in the head region of cingulin was implicated in the regulation of cingulin conformation and cytoskeletal interactions (Yano et al., 2013; Yano et al., 2018), we also generated combined dephospho-and phospho-mimetic mutants of the three relevant Ser residues in the head (Ser 137, 139, 154) of canine cingulin (Phospho Head, Fig. 2A, 2H and Dephospho Head, Fig. 2A, 2I).

**Figure 2.**
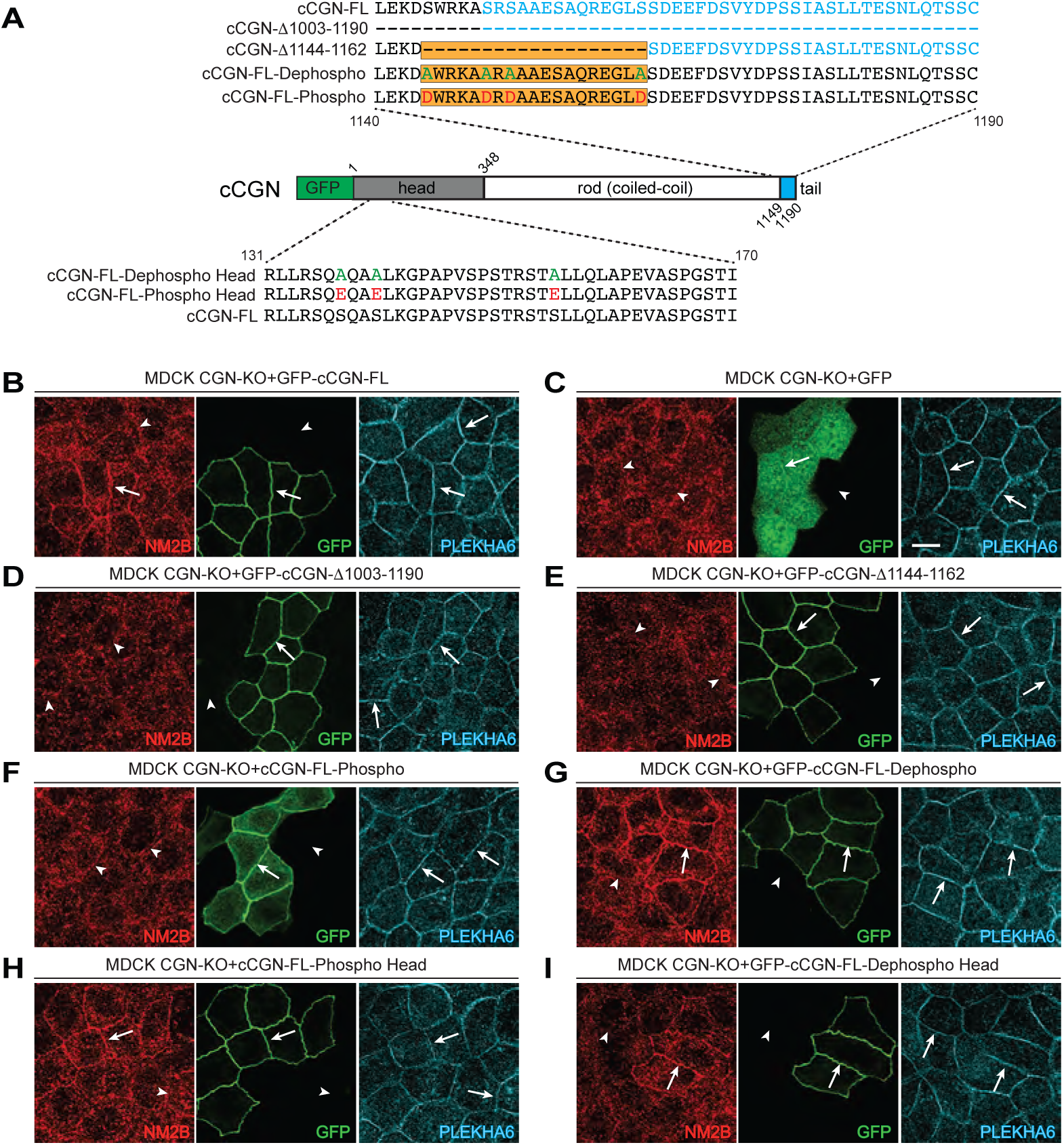
Phosphorylation of Ser residues within a 19-residue sequence in the rod-tail hinge, but not in the head domain of cingulin, regulates the junctional accumulation of NM2B. (A) Top: C-terminal sequences of WT canine CGN (cCGN-FL, region 1140–1190 with specific residues indicated in the region), or the truncation of the C-terminal rod-tail region of cingulin (cCGN-Δ1003-1190), or the truncation of 19-residue sequence in the rod-tail hinge (cCGN-Δ1144-1162), or mutant of hinge replaced with Ala residues (cCGN-1144-1162Ala), or where four Ser were mutated into Ala (cCGN-FL-dephospho-mimetic mutant), or Asp (cCGN-FL-phospho-mimetic mutant). Middle: scheme of GFP-tagged canine CGN (cCGN-FL), with the GFP tag (green, N-terminal), globular head (gray), coiled-coil rod (white) and globular tail (blue) domains (amino-acid residue boundaries are indicated above the scheme). Bottom: N-terminal sequences of WT canine CGN (cCGN-FL, region 131–170, with specific residues indicated in the region) and corresponding sequences of the cCGN-phospho-mimetic or-dephospho-mimetic head mutants. (B-I) IF microscopy analysis and localization of junctional labeling of NM2B in CGN-KO MDCK cells rescued with GFP-cCGN-FL (positive control, panel B), GFP–Myc alone (negative control, panel C) GFP-cCGN-Δ1003-1190 (D), GFP–cCGN-Δ1144-1162 (E), GFP-cCGN-Phos (F), GFP-cCGN-Dephos (G), GFP-cCGN-Phospho head (H), GFP-cCGN-Dephospho head (I). PLEKHA6 was used as a junctional reference marker. Arrows and arrowheads show normal and decreased/undetected junctional labeling, respectively. A representative image, derived from three independent experiments, is shown. Scale bar (C): 10 μm.

Using as an assay the immunofluorescent junctional localization of endogenous NM2B in the background of CGN-KO cells rescued with the different constructs, we confirmed that WT cingulin rescues junctional NM2B (arrows in Fig. 2B, PLEKHA 6 in blue as internal reference marker for junctions), whereas either GFP alone or C-terminally truncated CGN (Δ1003-1190) do not (negative controls, arrowheads in Fig. 2C-D) (Rouaud et al., 2023). Importantly, no rescue of junctional NM2B was observed upon either deletion of the hinge region (Δ1144-1162, arrowheads, Fig. 2E) or phospho-mutation of the 4 Ser within the hinge region (arrowheads, Fig. 2F). Instead, dephospho-mutation of the hinge region resulted in strong recruitment of NM2B to junctions (arrows, Fig. 2G). Finally, concerning the mutations of the Ser residues in the head domain of cingulin, neither phospho or dephospho mutation affected NM2B recruitment to junctions, that was indistinguishable from WT cingulin (arrows in Fig. 2H-I). In summary, phosphorylation of cingulin in the hinge region, but not in the head, regulates NM2B recruitment by cingulin to TJs in cells.

### Phosphorylation of the rod-tail hinge region of cingulin controls cingulin-NM2B interaction in vitro

To verify that the phosphorylation-mediated interaction between cingulin and NM2B detected in cells correlates with similar patterns of interactions in vitro, we used a GST pulldown assay. The purified, bacterially expressed cingulin-binding region of NM2B (GST-cNM2B-1757-2006, Fig. 3A, C, E, G) was used as a bait, and the GFP-tagged constructs of full-length WT and mutant canine cingulin were used as preys. Wild-type cingulin (GFP-cCGN-FL) was the positive control (Fig. 3A, C, G, E), and a truncation mutant of cingulin that does not bind to NM2B (GFP-cCGN-Δ1003-1190) was used as a negative control for NM2B binding (Fig. 3A) (Rouaud et al., 2023). As an additional control, we carried out parallel GST pulldowns using the C-terminal, cingulin-binding region of ZO-1 as a bait (GST-mZO-1-CterL, Fig. 3B, D, F, H). Immunoblot (IB) analysis of pulldowns using antibodies against GFP showed that deletion of the hinge alone (Δ1144-1162) abolished NM2B-cingulin interaction as effectively as the larger deletion (Δ1030-1190) (Fig. 3A). This indicated that the hinge region is the minimally required sequence for cingulin-NM2B interaction not only in cells, but also in vitro. In contrast, both deletion mutants interacted with ZO-1 as well as WT cingulin (Fig. 3B). Mutating all residues of the hinge region to Ala also abolished interaction of cingulin with NM2B, but not with ZO-1 (G-cCGN-1144-1162Ala, Fig. 3C-D). This indicated that the effect of the deletion of the hinge is not a consequence of artificial change in the length of the polypeptide chain. In agreement with the IF observations (Fig. 2F-G), the Ser->Ala dephospho-mimetic mutant (depho-1) of the hinge region (Fig. 2A) still interacted with NM2B (Dephospho in Fig. 3E), whereas the Ser->Asp phospho-mimetic mutant (pho-1) resulted in loss of binding to the NM2B bait (Phospho in Fig. 3E). Both dephospho and phospho mutants of the hinge interacted normally with the ZO-1 bait (Fig. 3F). Finally, both dephospho and phospho mutants of the head region bound similarly well to the NM2B (Fig. 3G) and the ZO-1 (Fig. 3H) baits, in agreement with the observations on cells (Fig. 2H-I). In summary, the hinge region is required for cingulin interaction with NM2B both in cells and in pulldown assays, whereas phosphorylation of Ser residues within this region negatively regulates cingulin-NM2B interaction both in cells and in vitro. Instead, phosphorylation of Ser residues either in the hinge or in the head does not affect the interaction of cingulin with ZO-1, consistent with the notion that interaction with ZO-1 only involves the ZIM region of cingulin (D’Atri et al., 2002; Vasileva et al., 2022). Moreover, either phosphorylation or dephosphorylation of Ser residues in the head does not affect binding of cingulin to NM2B.

**Figure 3.**
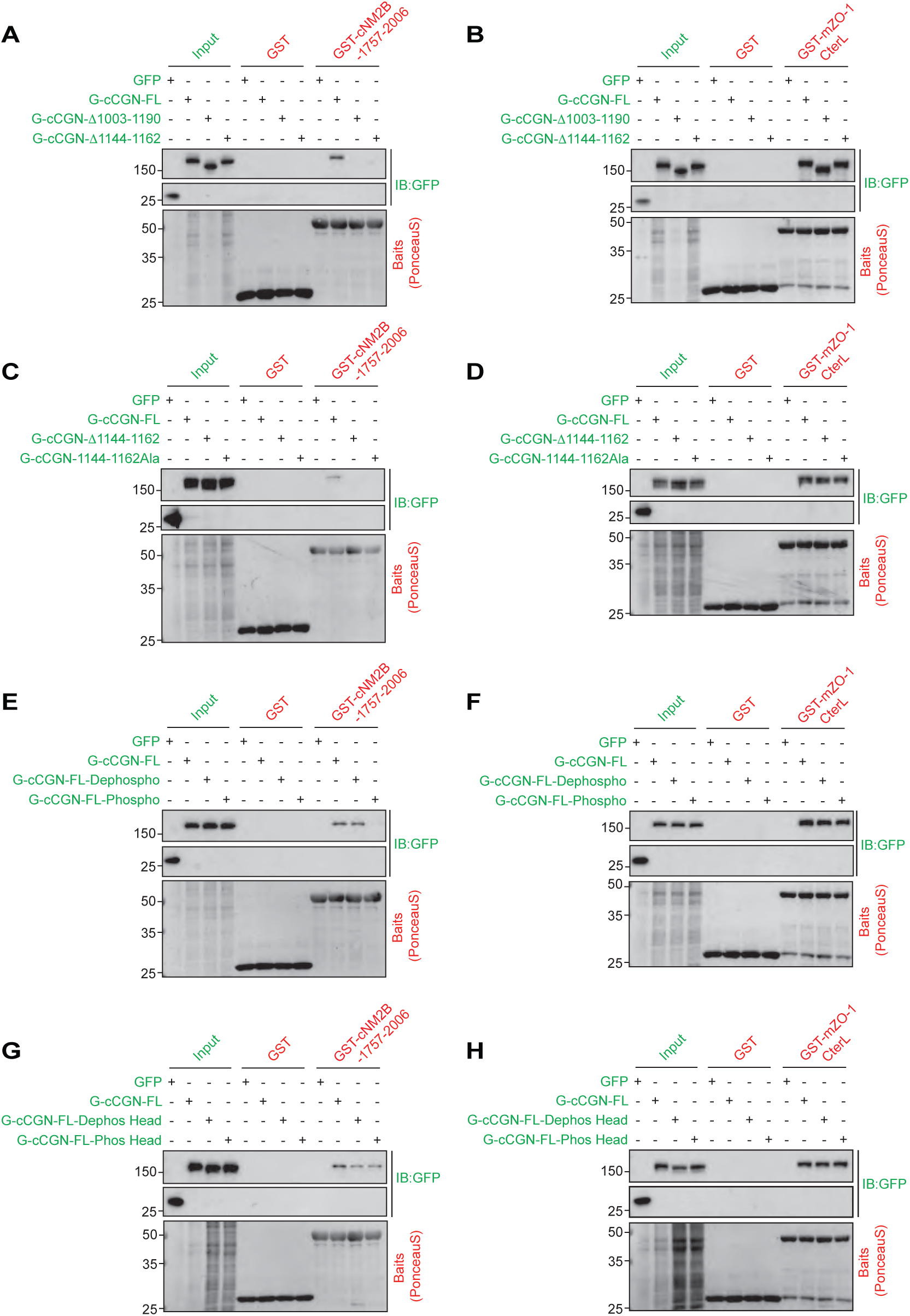
Phosphorylation of Ser residues in the hinge, but not within the head domain, controls interaction of cingulin with NM2B but not ZO-1 in vitro. (A-H) IB analysis using anti-GFP antibodies of pulldowns using GST-tagged affinity-purified fragments of either the last 250 residues of cNM2B (CGN-binding region, GST-cNM2B 1757–2006, Rouaud et al., 2023) (A, C, E and G) or mouse ZO-1 (CGN-binding region, GST-mZO-1CterL;Vasileva et al., 2022) (B, D, F and H) as baits. GST alone was the negative control bait. Preys were either GFP-myc (A-H, GFP, negative control) or GFP-tagged full-length CGN (A-H, G-cCGN-FL, positive control), or truncated CGN (Δ1003-1190) (A-B), or deleted hinge (Δ1144-1162) (A-D), or mutant of hinge replaced with Ala residues (cCGN-1144-1162Ala) (C-D), or phospho-or dephospho-mimetic of the hinge of CGN (E-F), or phospho-or dephospho-mimetic mutants of the head of CGN (G-H). Input lanes show normalized preys. Bottom panels show Ponceau Red-labeled baits. Numbers on the left indicate migration of pre-stained markers.

### Phosphorylation of Ser residues in the hinge, but not in the head domain, controls cingulin conformation in cells

Previous experiments in vitro using purified cingulin suggest that cingulin exists either in an open or a closed conformation, where the coiled-coil rod is either extended or folded, respectively, and that AMPK-dependent phosphorylation promotes the open conformation (Yano et al., 2018). Our previous studies suggest the conformation of cingulin in cells depends on interaction with NM2B (Rouaud et al., 2024), leading to the prediction that the cingulin hinge phospho-mutant will behave similarly to previously examined cingulin mutants that do not bind NM2B.

To test this hypothesis, mutant cingulin constructs with a N-terminal GFP tag and a C-terminal myc tag (green and red, Fig. 4) (Rouaud et al., 2024) were expressed in the background of CGN-KO MDCK cells. In this assay, the N-terminal cingulin tag (green, Fig. 4) co-localizes with ZO-2, used as a membrane-proximal TJ reference marker (blue, Fig. 4), and the C-terminal cingulin tag (red, Fig. 4) is either detected as two separate lines (extended conformation) or as one line (folded conformation), co-localizing with ZO-2 (Rouaud et al., 2024).

**Figure 4.**
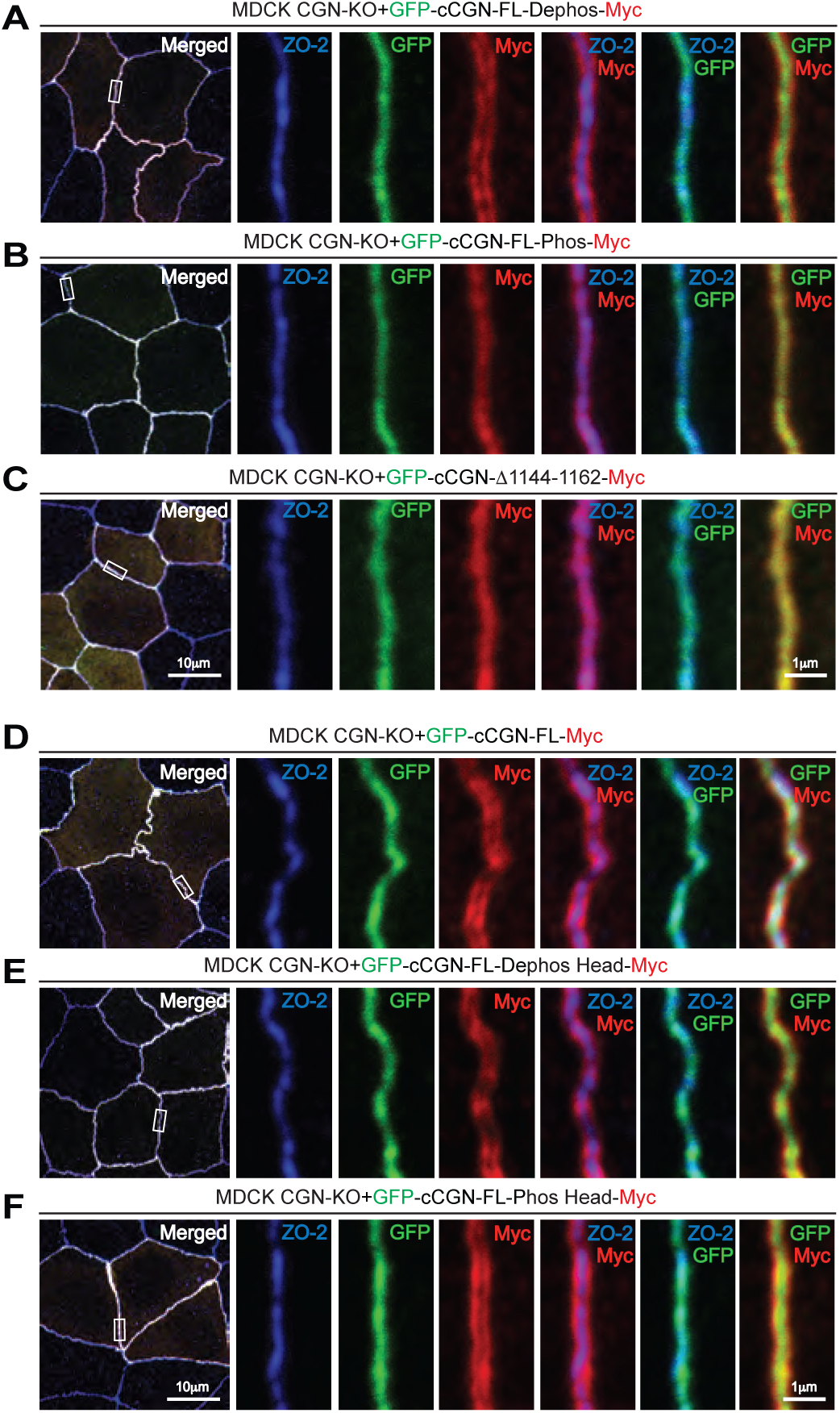
Phosphorylation of Ser residues in the hinge but not in the head domain controls cingulin conformation in cells (A-F) IF microscopy analysis of the localization of ZO-2 (blue, internal marker for TJ midline), and of the N-terminus (GFP, green) and C-terminus (Myc, red) of cingulin in CGN-KO MDCK cells rescued with cCGN-dephospho mutant (A), cCGN-phospho mutant (B), hinge truncation (cCGN-Δ1144–1162)(C), WT cingulin (GFP-cCGN-FL-Myc, positive control) (D), cCGN-Dephos Head mutant (E) or cCGN-Phos Head mutant (F) constructs. Collapse of the C-terminus of cingulin to the mid-zone (B-C) suggests a closed conformation of cingulin. High-magnification panels on the right correspond to highlighted white boxes in low-magnification micrographs on the left and show individual blue (ZO-2), green (GFP, cingulin N-terminus), red (myc, cingulin C-terminus) channels as well as merge images of bleu-red, blue-green, and red-green channels, for additional detail. One representative image from two independent experiments is shown. Scale bars: 10 μm (low magnification); 1 μm (high magnification).

In agreement with our prediction, cells expressing the hinge Dephospho mutant showed a clear separation between the myc-tagged C-termini of cingulin molecules on the two sides of the junction, indicating an open conformation (Fig. 4A, two separate lines in Myc red channel). Instead, both the Phospho mutant and the mutant lacking the hinge region (Δ1144-1162) showed only one line, suggesting a folded conformation (Fig. 4B-C, one central Myc-labelled line in red channel). Importantly, both Dephospho and Phospho mutants of the head domain behaved similarly to WT cingulin, because the C-termini of cingulin on the opposite sides of the junction were in all cases clearly resolved, and did not colocalize with ZO-2 (Fig. 4D for WT and Fig. 4E-F for head mutants).

Collectively, our results support the conclusion that binding of cingulin to NM2B is regulated by phosphorylation of Ser residues in the cingulin hinge, and is required for cingulin to maintain an open conformation in cells, whereas phosphorylation of head Ser residues does not affect either cingulin interaction with NM2B and ZO-1, junctional localization or conformation in cells.

### Phosphorylation of Ser residues in the rod-tail hinge but not in the head domain affect TJ membrane tortuosity in cells

TJ membrane tortuosity is a readout of the balance between perpendicular and parallel forces exerted on the TJ membrane, and depends on the organization and contractility of the junctional actomyosin cytoskeleton (Choi et al., 2016; Mauperin et al., 2025b; Rouaud et al., 2023; Tokuda et al., 2014; Van Itallie et al., 2009) (reviewed in (Lynn et al., 2020; Tang, 2018). Since AMPK-dependent phosphorylation of Ser residues in the cingulin head was reported to regulate its interaction with actin and microtubules in vitro, we asked whether phosphorylation of either the head or the hinge could affect TJ membrane tortuosity, by regulating cingulin interaction with the actomyosin cytoskeleton. To address this question, we expressed wild-type and mutant cingulin constructs in the background of CGN-KO cells and examined the tortuosity of TJ using ZO-2 as a membrane-proximal marker (Fig. 5).

**Figure 5.**
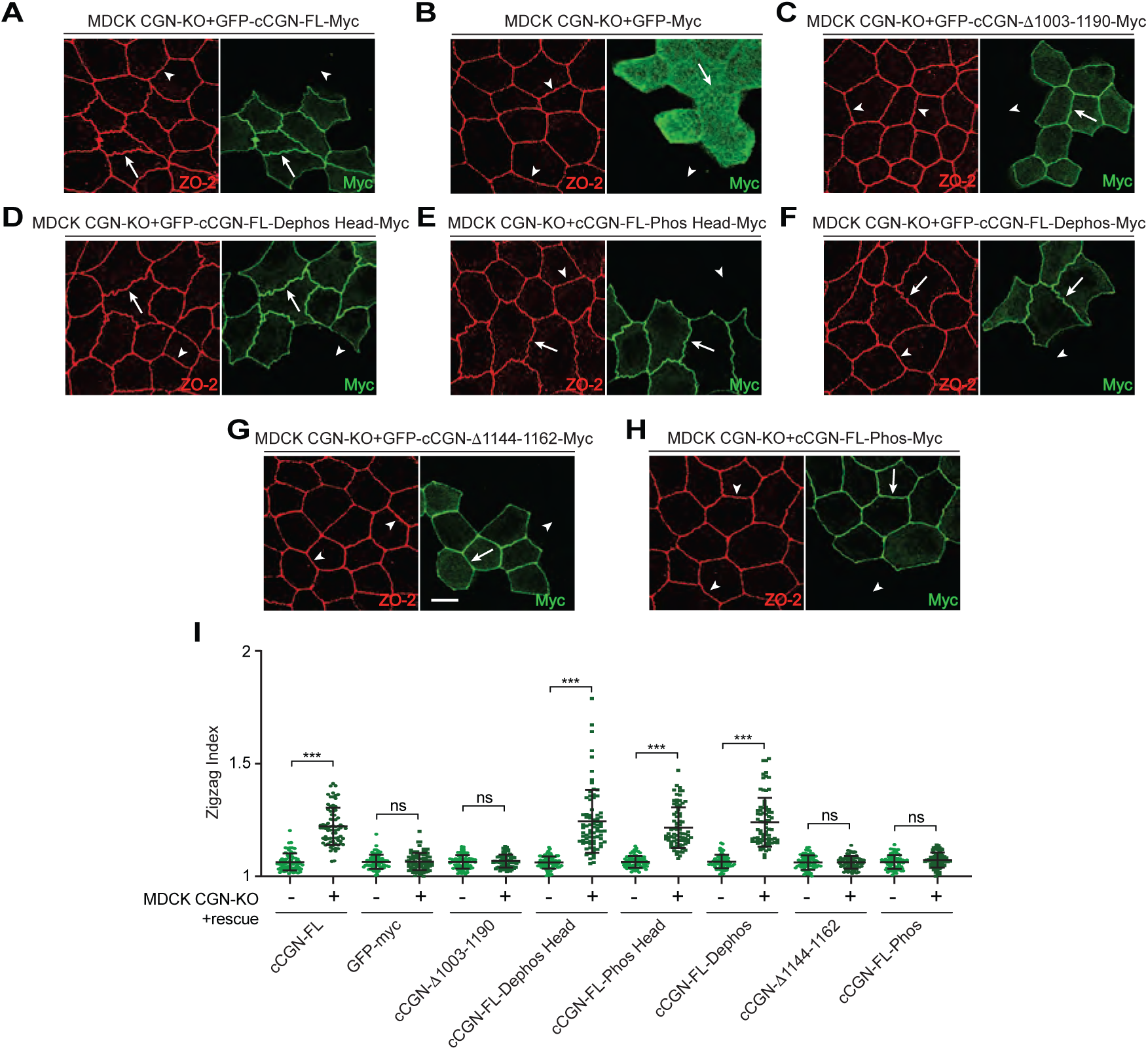
Phosphorylation of Ser residues of cingulin within the hinge, but not in the head domain, affect TJ membrane tortuosity in cells (A-I) IF microscopy analysis and quantification of the tortuosity for CGN-KO cells rescued with either full-length GFP-myc-tagged canine CGN (GFP–cCGN-FL-Myc) (A), or GFP–Myc alone (negative control) (B), GFP-cCGN-Δ1003-1190 (C), or CGN-Dephospho head (D), or cCGN-Phospho head (E), or cCGN-Dephospho (F), or truncation of the hinge (GFP–cCGN-Δ1144-1162) (G), or cCGN-Phos (H). Arrows and arrowheads show junctions with normal and decreased tortuosity, respectively. A representative image, derived from two independent experiments, is shown. Scale bar: 10 μm. In (I) (-) and (+) refer to the same cell line without and with exogenous expression of the rescue construct. Dots show replicates (n=75) and bars represent mean ± SD. Statistical significance was determined by one way ANOVA test, ***p≤0.001.

As shown previously, CGN-KO cells showed decreased TJ membrane tortuosity, and this phenotype was rescued by expression of wild-type cingulin (Fig. 5A, quantifications in Fig. 5I) (Rouaud et al., 2023). In contrast, expression of either GFP (negative control, Fig. 5B, quantifications in Fig. 5I), or of a mutant of cingulin lacking the last 187 residues, and that does not bind to NM2B, did not rescue tortuosity (Fig. 5C, quantifications in Fig. 5I) (Rouaud et al., 2023). Importantly, TJ tortuosity was rescued by expression of cingulin with either dephospho or phospho mutations in the head (Fig. 5D and 5E, quantifications in Fig. 5I), and with dephospho mutations of the hinge (Fig. 5F, quantifications in Fig. 5I). Instead, tortuosity was not rescued by either deletion of the 19-residue hinge (Fig. 5G, quantifications in Fig. 5I), or by expression of the phospho-hinge mutant (Fig. 5H, quantifications in Fig. 5I).

Together, these results indicate a strict correlation between binding of cingulin to NM2B in vitro and in cells and its ability to rescue TJ membrane tortuosity in cells.

### Protein kinases CK2 and CK1 are involved in the phosphorylation of cingulin in vitro and in cells

To analyze further the mechanism of cingulin phosphorylation, we sought to identify protein kinases which might be involved. Two residues of the hinge region, namely Ser1162 and Ser1163, show a consensus sequence for phosphorylation by protein kinase CK2 (Salvi et al. 2009; Meggio e Pinna 2003). CK2 is a ubiquitously expressed, constitutively active kinase, that phosphorylates hundreds of substrates and is involved in multiple diseases (Borgo, D’Amore, Sarno, et al. 2021). To assess CK2 ability to phosphorylate these cingulin residues, we performed in vitro kinase assays with recombinant purified CK2, using as a substrate a bacterially expressed, purified recombinant fragment of the rod-tail region of cingulin. This fragment was produced either as WT (residues 1015-1203 of the human sequence, where Ser1175 and Ser1176 correspond to Ser1162 and Ser1163 of the canine sequence), or as variants where Ser1175 and Ser1176 were replaced by alanine (either individually or both).

The WT C-terminal cingulin fragment was phosphorylated by CK2 (GST-hCGN WT in Fig. 6A, Ponceau red staining of protein loadings in Fig. 6B). In contrast, mutation of either Ser1175 or Ser1176 into Ala resulted in significant decrease in phosphorylation (Fig. 6A-B, quantifications in Fig. 6C). Significantly, simultaneous mutation of both and Ser1175 and Ser1176 almost completely abolished phosphorylation, indicating that these two residues are major phosphorylation sites by CK2 (Fig. 6A-C).

**Figure 6.**
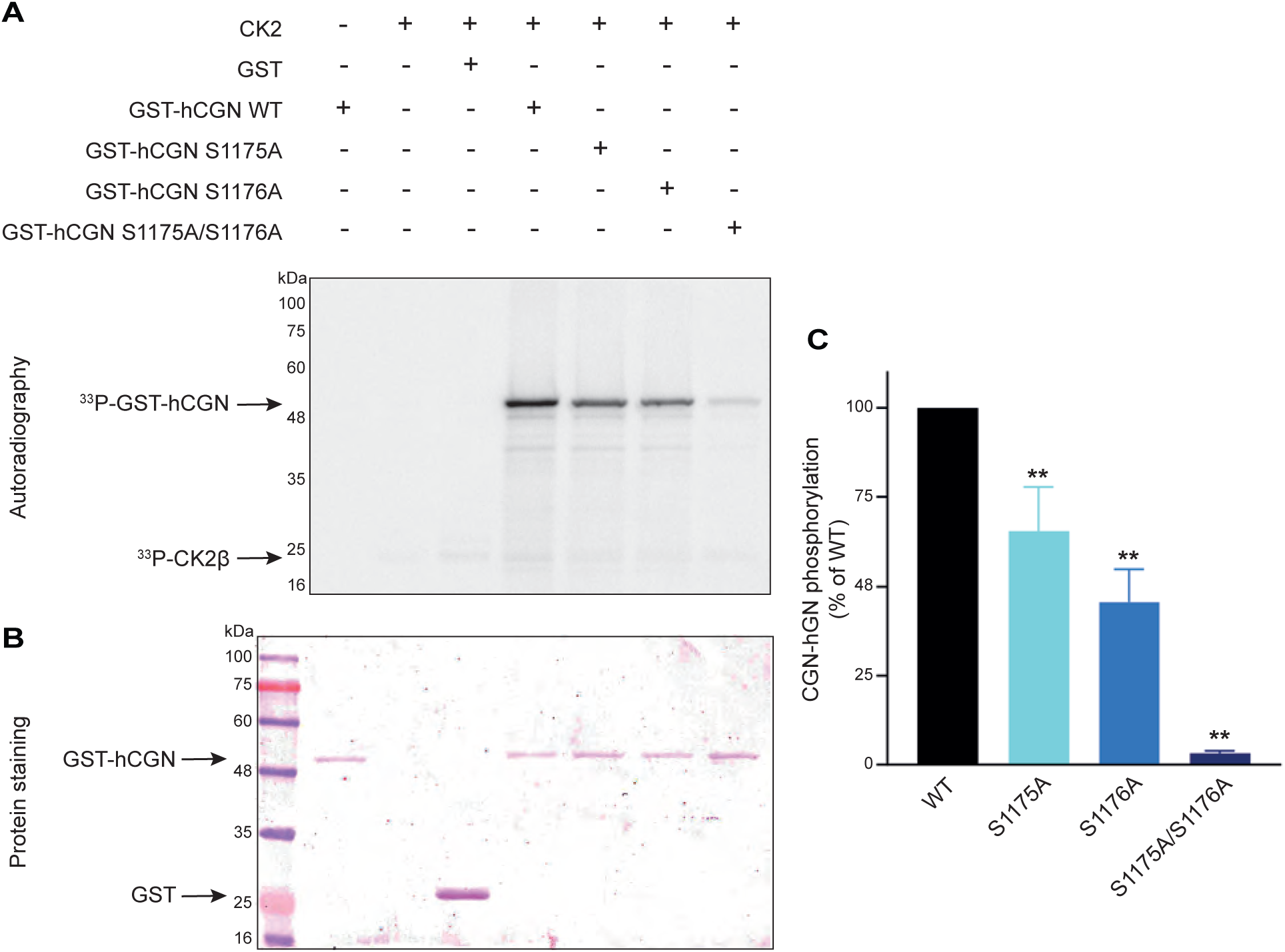
Protein kinase CK2 phosphorylates Ser1175 and Ser1176 of the human cingulin hinge in vitro. (A-B) Autoradiogram (A) and corresponding Ponceau-S protein staining (B) of SDS-PAGE analysis of blots of phosphorylation reactions. Recombinant CK2 and the indicated purified recombinant substrates indicated on the top lines (GST alone, negative control, and GST-CGN WT and mutant fragments) were incubated in a radioactive [γ-^33^P]-ATP-containing phosphorylation mixture (see Methods). Samples were separated by SDS-PAGE followed by blotting on a PVDF membrane. (C) Quantification of phosphorylation of the indicated GST-tagged cingulin recombinant proteins, as percentage compared to the WT control, taken as 100% (means +/-SD of three independent experiments). ** p<0.01.

To examine the physiological relevance of this phosphorylation, we focused on Ser1175, corresponding to Ser1162 in canis, which shows a consensus phosphorylation sequence for both CK2 and CK1 (Venerando et al., 2014). We therefore used chemical inhibitors of CK2 kinase (CX4945) (Pierre et al., 2011) and CK1 kinase (D4476) (Rena et al., 2004) to probe their roles in cultured cells (Fig. 7). Full-length WT cingulin rescued NM2B accumulation at junctions either in the absence (Fig. 7B) or in the presence of either the CK1 or the CK2 inhibitor (Fig. 7D-F), consistent with the concept that dephosphorylation promotes cingulin-NM2B interaction. Next, we focused on the depho-6 mutant of cingulin, where all the Ser residues of the hinge are mutated to Ala, except for Ser 1162 (Fig. 7A). The depho-6 mutant did not rescue junctional NM2B in CGN-KO cells (Fig. 7C, see also Fig. 1K’), suggesting that allowing phosphorylation at this site prevents cingulin-NM2B interaction. Strikingly, in the presence of either the CK1 or the CK2 inhibitors, the depho-6 mutant of cingulin became capable to rescue the junctional accumulation of NM2B (Fig. 7E and 7G, respectively). This observation suggests that in cells phosphorylation of Ser1162 by either CK1 or CK2 negatively regulates cingulin-NM2B interaction.

**Figure 7.**
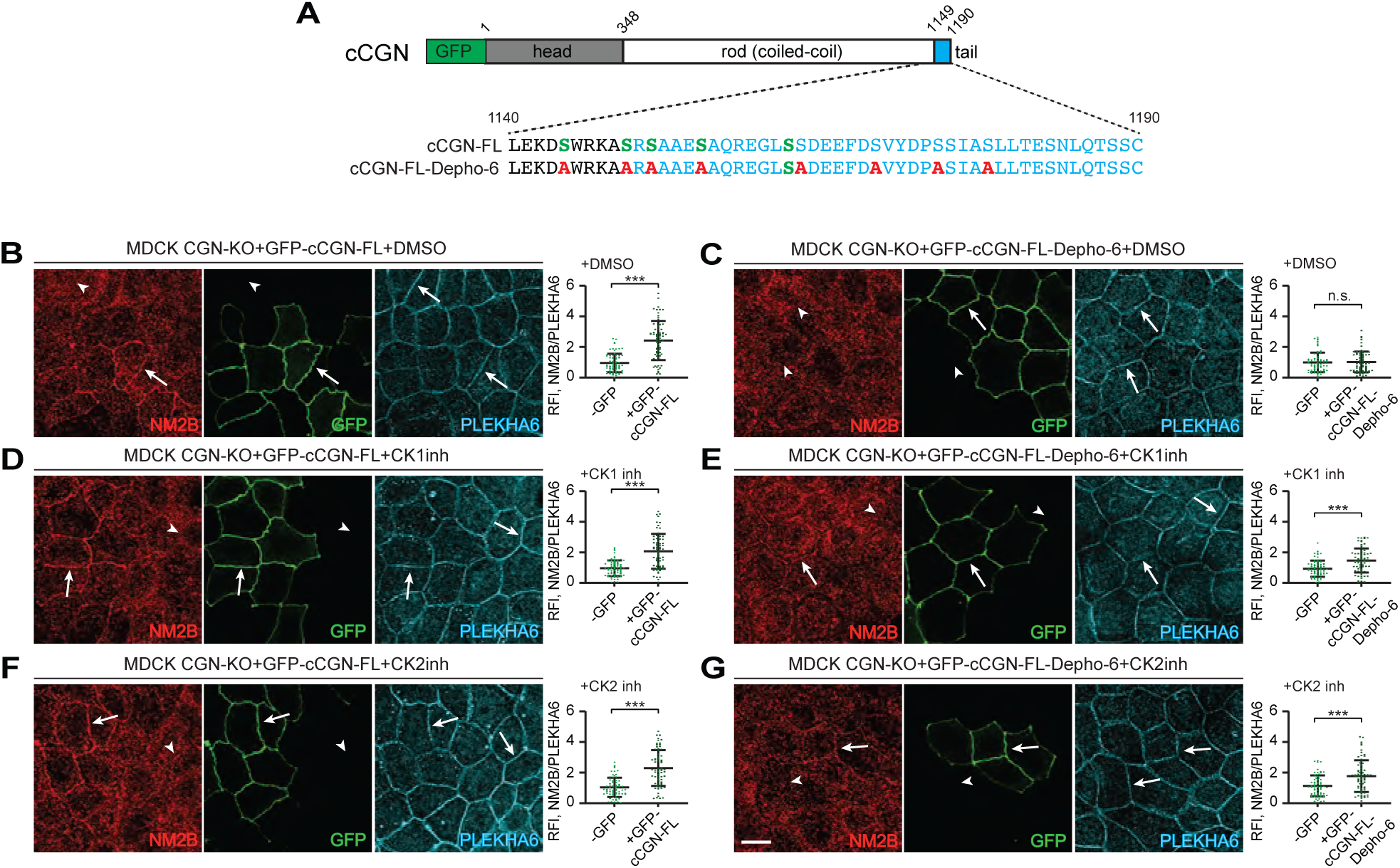
Inhibition of either CK1 or CK2 promotes the rescue of junctional NM2B in cingulin-KO cells expressing the depho-6 mutant of cingulin. (A) Top: scheme of GFP-tagged canine CGN (cCGN-FL), with the GFP tag (green), globular head (gray), coiled-coil rod (white) and globular tail (blue) domains (amino-acid residue boundaries are indicated above the scheme). Bottom: C-terminal sequences of WT canine CGN (cCGN-FL, sequence 1140–1190 with specific residues indicated in the region) and corresponding sequences of the cCGN-dephosphomimetic-6 mutant (depho-6), which does not rescue junctional NM2B (Fig. S1K’). (B-G) IF microscopy analysis and localization (left) and quantification of junctional labeling (relative fluorescence intensity) (right) of NM2B in CGN-KO MDCK cells rescued with GFP-cCGN-FL (B, D and F), or with GFP-cCGN-depho-6 mutant (C, E and G) treated either with DMSO (B and C) or with CK1 inhibitor (D and E: 25 μM, 8h) or with CK2 inhibitor (F and G: 25 μM, 14h). Arrows and arrowheads show increased, normal and decreased/undetected junctional labeling, respectively. Quantifications of relative fluorescent intensity (RFI) shows the ratio between the junctional staining of NM2B versus the junctional marker PLEKHA6 (n=70 junctions) from three independent experiments. Data in quantifications are represented as mean±SD. Statistical significance was determined by unpaired Mann-Whitney’s test. ***p≤0.001. Scale bar (G)= 10 μm.

## DISCUSSION

The actomyosin cytoskeleton associated with the apical junctional complex is critically important for junction integrity and TJ barrier function, and several studies have described a role for protein kinases such as ROCK kinase, protein kinase C (PKC) and myosin light chain kinase (MLCK) in TJ assembly and disassembly, regulation of barrier function, and the organization and contractility of the junctional actomyosin cytoskeleton (Clayburgh et al., 2004; Graham et al., 2019; He et al., 2020; Nusrat et al., 1995; Samarin et al., 2007; Suzuki et al., 2009; Turner et al., 1997). However, little is known about the role of phosphorylation of specific TJ proteins in regulating their dynamic interaction with actin, microtubules and cytoskeletal motors. This study provides new evidence that the function of cingulin as a tether for NM2B at TJs is regulated by phosphorylation. In addition to refining the mapping of the sequences required for NM2B binding to a 19-residue sequence at the hinge between coiled-coil rod and globular tail of cingulin, we confirm that loss of binding to NM2B, either by deletion or phosphorylation of the NM2B-binding region, results in altered cingulin conformation and loss of TJ membrane tortuosity. This supports the idea that the integrity of the ZO-1-cingulin-NM2B module is crucial for the mechanical properties of the TJ membrane cortex (Rouaud et al., 2023; Rouaud et al., 2024). Finally, we identify relevant phosphorylation sites in the hinge and provide evidence that protein kinases CK1 and CK2 may be involved in cingulin phosphorylation.

Our experiments show that cingulin phosphorylation in the hinge regulates cingulin-NM2B interaction. This type of regulation has been demonstrated for NM2s, where phosphorylation of the C-terminal rod-tail hinge stabilizes the folded conformation of myosin and inhibits rod-rod interaction and filament assembly, by introducing localized negative electrostatic charges in the tail, that disrupt the parallel and antiparallel packing of the rods (Dulyaninova and Bresnick, 2013; Rosenberg et al., 2008; Vicente-Manzanares et al., 2009). Our results suggest that a similar charge-dependent mechanism regulates the interaction between cingulin rod-tail and NM2B rod-tail. On the other hand, we show that phosphorylation of conserved Ser residues in the cingulin head affects neither interaction with NM2B, nor cingulin conformation, nor TJ membrane tortuosity. This suggests that the previously observed effects of AMPK-dependent phosphorylation on cingulin conformation in vitro (Yano et al., 2018) are not relevant at junctions in cells. However, head phosphorylation by AMPK could still affect interaction with microtubules (Yano et al., 2013) and perhaps participate in modulating the transition between folded (closed) and extended (open) state and cytoskeletal interactions of the soluble, cytoplasmic fraction of cingulin, which exchanges with a junction-associated fraction (Paschoud et al., 2011). Thus, the exchange between soluble and the junction-associated cingulin fractions may depend both on the presence of ZO-1 and NM2B ligands in the neighboring TJ and cytoskeletal cortex, respectively, and on cingulin head phosphorylation. In this model, interaction with ZO-1 and NM2B and dephosphorylation of cingulin would be required to promote transition of cingulin conformation to an open state at TJs, and its stabilization. These hypotheses should be tested in the future.

The physiological relevance of cingulin phosphorylation remains to be established. Several post-translational modifications of TJ transmembrane proteins affect barrier function (Reiche and Huber, 2020). Although there is no clear evidence so far that KO of cingulin has an impact on epithelial barrier function in vitro and in vivo (Guillemot et al., 2004; Guillemot et al., 2012; Mauperin et al., 2023), the loss of cingulin affects endothelial permeability (Schossleitner et al., 2016; Zhuravleva et al., 2020). Thus, it cannot be ruled out that cingulin phosphorylation affects barrier function in certain tissues and physiological and pathological states. If this is the case, it appears unlikely that the mechanism is through regulation of interaction with NM2B, because NM2B-KD/KO cells show normal TJ barrier function (Ivanov et al., 2007; Mauperin et al., 2025a), and in mice genetic ablation of NM2B results in embryonic lethality due to cardiac and neuronal defects, with no reported epithelial barrier phenotype (Tullio et al., 1997; Tullio et al., 2001). Moreover, paracingulin-KO cells and mice do not display detectable defects on barrier function of epithelial and endothelial tissues at steady-state, despite decreased NM2 association with junctions (Mauperin et al., 2023; Rouaud et al., 2025a; Rouaud et al., 2025b). Therefore, it appears unlikely that phosphorylation-dependent regulation of NM2B tethering by either cingulin or paracingulin might significantly affect barrier function, and additional studies are required to clarify these questions.

One possible physiological role of cingulin hinge phosphorylation/dephosphorylation could be during junction assembly and disassembly that occurs during epithelial cell polarization and cell division, respectively. Cyclin-dependent kinase-and mitogen-activated kinase-centered interaction networks are coordinately down-and up-regulated in late mitosis, respectively (Dulla et al., 2010), and downstream activation of kinases that phosphorylate cingulin could occur at different steps of mitosis. In this respect, our findings, suggesting a role for CK2 and CK1 in cingulin phosphorylation may seem counterintuitive, since these kinases are considered constitutively active, and do not depend on second messengers or other cellular events to become catalytically competent. However, the activity of these kinases can be modified under certain cellular circumstances. For example, CK2 has essential roles in mitotic progression (Canton and Litchfield, 2006; D’Amore et al., 2019; St-Denis et al., 2011), is implicated in the regulation of cell morphology and cytoskeletal structures, thus affecting cell shape, movement and division (Borgo et al., 2021a; Borgo et al., 2021b; Venerando et al., 2014). For example, CK2 has essential roles in mitotic progression (Canton and Litchfield, 2006; D’Amore et al., 2019; St-Denis et al., 2011), is implicated in the regulation of cell morphology and cytoskeleton structures, thus affecting cell shape, movement and division (Canton and Litchfield, 2006; D’Amore et al., 2019), and directly regulates microtubule dynamics (Lim et al., 2004). CK2 has also been reported to regulate TJs, by affecting TJ protein complex formation (Cordenonsi et al., 1999b; Dorfel et al., 2013; McKenzie et al., 2006; Raleigh et al., 2011). Similarly, CK1 family enzymes have major regulatory functions in microtubule and cytoskeletal dynamic processes (Roth et al., 2022), and interacts with cingulin, based on mass spectrometry studies (Boldt et al., 2016). Thus, although our observations suggest that CK1 and CK2 are relevant for the regulation of NM2B/cingulin interaction, additional studies will be required to establish their respective relevance and relative importance with respect to other redundant kinases that may phosphorylate cingulin. Moreover, since here we used a pan-CK1 inhibitor (Rena et al., 2004) it remains to be determined which CK1 isoform(s) is involved. Finally, it remains to be clarified whether phosphorylation of paracingulin regulates its interaction with NM2s, and whether NM2B rod/tail phosphorylation regulates in turn its interaction with cingulin and paracingulin.

In summary, our study refines the structural requirements of cingulin-NM2B interaction by identifying a 19-residue hinge region that is required for this interaction. We show that phosphorylation of Ser residues in this region, but not in the head domain, controls the recruitment of NM2B by cingulin and downstream open conformation of cingulin and TJ membrane tortuosity in cells. Finally, we provide evidence for a potential role of protein kinases CK1 and CK2 in cingulin phosphorylation.

## METHODS

### Experimental models and transfection

Culture and transfection of MDCK (Madin-Darby Canine Kidney II cell line, female) and HEK293T cells was as described in (Rouaud et al., 2023). Drugs treatments were as follows (final concentration, duration, catalog number and source): CK1 inhibitor (25 μM, 8 h, D4476 MedChemExpress) and CK2 inhibitor (5 μM, 14 h, CX4945 Selleckchem).

### Antibodies and plasmids

The following primary antibodies against the indicated proteins were used for either immunoblotting (IB) or immunofluorescence microscopy (IF) at the indicated dilutions: mouse cingulin (22BD5A1; IF: 1:500); rabbit NM2B (909901; IF: 1:250); rat PLEKHA6 (RtSZR127; IF: 1:100); mouse GFP (11814460001; IF: 1:200, IB: 1:1000); goat ZO-2 (sc-8148; IF: 1:100); rabbit myc (06-549; IF: 1:200). The specificity of the antibodies was established previously using IB and IF on KO/KD cells and tissues (Rouaud et al., 2023). Secondary antibodies for immunoblotting were from Dako and diluted at 1:3000: polyclonal goat anti-mouse HRP (P0447). Secondary antibodies for immunofluorescence were from Jackson ImmunoResearch (1:300): anti-mouse-IgG (715-546-151)-Alexa Fluor 488; anti-rabbit-IgG (711-165-152)-Cy3; anti-rat-IgG (712-175-153)-Cy5; anti-goat-IgG (705-606-147)-Alexa Fluor 647.

Constructs of GFP-myc-his in pCDNA3.1(−) (Guerrera et al., 2016), and GFP-cCGN-FL-myc, GFP-cCGN-Δ1003-1190-myc in pCDNA3.1(-) (Rouaud et al., 2023) were described previously. The new constructs were generated by PCR amplification with appropriate oligonucleotides and subcloned into the indicated cloning sites (numbers in parentheses indicate construct numbers in the Citilab collection): GFP-cCGN-FL-myc-phosphomimetic (S3020 and S2872) and dephosphomimetic (S3019 and S2871) and truncated mutant of CGN corresponding to 19-residue stretch in the rod-tail hinge (Δ 1144-1162) (S3002), ClaI-SbFI in pCDNA3.1(-). Region1144-1162 replaced by alanine amino acids (1144-1162A) (S3053) SbFI-BSSHII in pCDNA3.1(-). GFP-cCGN-FL-myc-phosphomimetic-2,-3,-4,-5,-6,-7,-8,-9,-10 (S2924, S2912, S2936, S2942, S2915, S2934, S2935, S2913 and S2914) and dephosphomimetic-2,-3,-4,-5,-6,-7,-8,-9,-10 (S2939, S2937, S2940, S2925, S2923, S2941, S2916, S2917 and S2938), SbFI-BssHII in pCDNA3.1(-). GFP-cCGN-FL-myc-phosphomimetic head and-dephosphomimetic head (S2377 and S2376), NotI-ClaI in pTRE2hyg. A schematic representation of the mutants is shown in Fig. 1A and Fig. 2A. Constructs of GST-tagged mZO1-C-terminal large (Vasileva et al., 2022) and GST-tagged last part of cNM2B (Rouaud et al., 2023) were described previously.

### Glutathione S-Transferase (GST) pulldown, SDS-PAGE and immunoblotting

For GST pulldowns, GST-tagged protein baits were expressed in BL21 bacteria and purified by affinity chromatography on magnetic beads as described in (Sluysmans et al., 2021). Preys were tagged with GFP and consisted of full-length and mutant or truncated proteins expressed in HEK293T cells. Preparation of lysates and pulldowns were as described in (Sluysmans et al., 2021).

SDS-PAGE electrophoresis, and immunoblotting was carried out as described previously (Rouaud et al., 2024; Sluysmans et al., 2021). Numbers on the left of IB in the Figures indicate migration of pre-stained molecular size markers (kD).

### Immunofluorescence microscopy and analysis of immunofluorescence data

Immunofluorescent labeling of cells on coverslips, confocal microscopy and preparation of Figures from images were as described in (Rouaud et al., 2024).

For the measurement of the zig-zag index (L(TJ)/L(St)) (ratio between actual length of bicellular junction and the distance between two vertexes), we used the method described in (Tokuda et al., 2014), and measured the length of the TJ (L(TJ)) by using the freehand line trace in ImageJ, and the straight length of junction (L(St)) by using a straight line between vertexes. Typically, 75 bicellular junctions were analyzed for each of independent duplicate experiments.

For the quantification of junctional immunofluorescent signal pixel intensity for each channel was measured in the selected junctional area using the polyhedral tool of ImageJ, and the averaged background signal of the image was subtracted. Relative intensity signal was expressed as a ratio between NM2B and an internal junctional reference (PLEKHA6). Typically, 70 junctional segments were analyzed, for each of independent triplicate experiments.

### In vitro phosphorylation of cingulin recombinant fragments

Purified recombinant α_2_β_2_ CK2 (10 ng) (Venerando et al., 2013) was incubated with 0.5 mg substrates (bacterially expressed, GST-tagged cingulin fragments purified by affinity chromatography) in a phosphorylation mixture containing 100 mM Tris–HCl pH 7.5, 10 mM MgCl_2_, 20 μM [γ-^33^P], ATP (1000–2000 cpm/pmol) and 0.1 M NaCl, in a final volume of 20 μl. The reaction was performed for 10 min at 30°C and stopped by addition of Laemmli buffer. Samples were separated on 11% SDS-PAGE, blotted on PVDF membrane, and analyzed for radioactive incorporation of ^33^P by CyclonePlus Storage Phosphor System, PerkinElmer. Ponceau staining of the membrane was used to assess protein loading.

### Quantifications and statistical analysis

Data processing and analysis were performed in GraphPad Prism 8. All experiments were carried out at least in duplicate, and data are shown either as dot-plots (with mean and standard deviation indicated). Statistical significance was determined by unpaired Mann-Whitney’s test (when comparing two sets of data), or one way ANOVA test and the normality distribution was tested with Shapiro-Wilk test (ns=not significant difference, and **** p≤0.0001).

